# Ecological interactions between host, commensal and pathogenic bacteria in models for the intestinal epithelium

**DOI:** 10.1101/2023.11.09.565308

**Authors:** Nening M. Nanlohy, Nina Johannesson, Lucas Wijnands, Laura Arroyo, Jelle de Wit, Gerco den Hartog, Katja C. Wolthers, Adithya Sridhar, Susana Fuentes

## Abstract

Gut host physiology and the microbiome intricately interact in the complex ecosystem of the human digestive tract, playing a crucial role in maintaining overall health. In recent decades, the role of the gut microbiota in the defence against pathogens and modulating local and distal immunity has been well-established. The interactions between commensal and potential pathogenic bacteria with the intestinal epithelium can initiate immune responses in the epithelial cells, which, in turn, activate downstream immune responses in other immune cells. These intricate processes involved, especially when multiple microorganisms are present as seen in the intestinal microbiome, remain only partially understood. Previously, it was observed that in adults aged 60 years or older, the commensal *Ruminococcus torques* (Rt) and *Escherichia coli* were associated with influenza-like illness and a heightened pro-inflammatory immune profile. In this study, we used a CaCo-2 cell-based model and a human intestinal enteroid (HIE) model to explore epithelial responses to Rt and an adherent invasive *E. coli* (AIEC) both individually and in co-cultures under anaerobic conditions. Additionally, CaCo-2 cells were co-cultured with peripheral blood mononuclear cells, revealing downstream activation of immune cells. While both systems showed comparable cytokine profiles, they differed in their responses to the different bacteria, with the organoid system being more representative for intestinal epithelial cells in humans. We provide mechanistic evidence of the pro-inflammatory responses associated with these bacteria in the intestinal ecosystem. These models, particularly in the context of combined infections, represent a valuable and promising avenue for future research. They contribute to a deeper understanding of the complex interactions between the gut microbiota, epithelial intestinal cells and immune cells in the gut ecosystem, thereby promoting advances in the field of gut health and host response.

## Introduction

Interactions between commensal microbes, host epithelium and the immune system in the gut mucosal environment contributes to maintenance of health [1]. This gut ecosystem plays an important role in the regulation of metabolic, endocrine and immune functions, including the defence against infections [2, 3]. Throughout life, the gut microbiota continuously evolves, influenced mainly by environmental and lifestyle factors [4]. Disturbances in the composition of the gut microbiota can impact the integrity of the gut barrier and are linked with the development of (chronic) inflammatory and metabolic diseases [5-7]. Furthermore, the proinflammatory state associated with aging, often referred to as inflammaging, has been associated with changes in the gut microbiome [8-10]. Recently, we showed that high abundance of the gut commensals *Ruminococcus torques* and *Escherichia coli*, often observed in inflammatory processes such as inflammatory bowel disease [11-13], were associated with a pro-inflammatory immune profile in the ageing population with influenza-like illness (ILI) [14]. Such observational studies can provide new perspectives but are often confounded by a multitude of factors, especially in the ageing population due to increased prevalence of comorbidities and medication use, and therefore, caution is warranted when making causal inferences.

To move beyond associations, more controlled mechanistic studies are key to obtain causal as well as deeper insights into the immunomodulatory effects of bacteria on the gut ecosystem. These should account for the trilateral interactions between human gut microbiota, epithelial intestinal cells and immune cells in the gut microenvironment [15, 16]. Currently, different in vitro models are available to study intestinal epithelial infections with varying levels of complexity and translatability [17]. Transwell co-cultures based on Caco-2 cell lines have been widely used to study intestinal host-microbe interactions [18, 19]. The CaCo-2 monolayer, originated from a colon adenocarcinoma, can spontaneously differentiate to resemble a small intestine-like phenotype and thereby, mimics important properties of the intestinal epithelium [18, 20]. These cultures can also be combined with peripheral blood mononuclear cells (PBMCs) to assess immune responses not only from the epithelial cells but also from immune cells mimicking the gut-associated lymphoid tissue. However, a gut epithelial model based on CaCo-2 cells lacks the degree of differentiation of epithelial cell subtypes present in vivo [16]. Recently developed human primary intestinal organoid models can alleviate some of these issues by recapitulating the in vivo cellular heterogeneity [21-26]. 2D intestinal epithelial monolayers, derived from primary human epithelial cells on Transwell*^®^* inserts [22, 25], facilitate infection on the luminal side and enable studying complex multicellular responses as well as introducing biological variability. However, the high costs of these models are a drawback that limits experimental scale-up and breadth [27].

In our study, we aimed to expand on our previous association study using these two aforementioned model systems to investigate interactions between key players in the gut ecosystem. We investigated the immune processes that occur in the gut epithelium primed with a commensal (*R. torques*) and subsequently challenged with a pathogenic (adherent invasive *E. coli*, AIEC) bacteria, selected for their previously observed synergistic interactions. We analysed the individual and combined impact of these bacteria in the cytokine and chemokine response by both gut epithelial models and PBMCs. Our work provides mechanistic evidence on the pro-inflammatory profile associated with these bacteria in the intestinal ecosystem.

## Materials & Methods

### Bacterial cultures and CaCo-2 cell-based model

Bacterial cultures of *Ruminococcus torques* (ATCC27756) and an adherent/invasive *Escherichia coli* (AIEC strain LF82), and CaCo-2 cells were prepared as previously described [14]. For in vitro experiments, *R. torques* were cultured overnight at 37°C under anaerobic conditions (Anoxomat, Hettich Instruments) and AIEC at 37°C under aerobic conditions. Bacteria were inoculated on the apical (AP) side of the epithelial monolayer at approximately 10e9 CFU/mL. Enumeration of unbound bacteria was performed after plating 10-fold dilutions of the Triton-extracts (data not shown). Columbia agar with sheep blood was used for determining *R.torques* and *AIEC.* In addition, Tryptone Bile X-Glucuronide (TBX) plates were used for *AIEC*.

### Human Intestinal Enteroid Monolayer Culture

#### Ethics statement

Human foetal intestinal tissues (gestational age 14-19 weeks) were obtained from the HIS Mouse Facility of the Amsterdam UMC through a private clinic. All material were collected from donors from whom a written informed consent for the use of the material for research purposes had been obtained by the clinic. These informed consents are kept together with the medical record of the donor by the clinic and information provided (age of the donor and gestational age of the foetus) to the Amsterdam UMC does not allow for identification of the donor without disproportionate efforts. The use of the foetal material and anonymized material for medical research purposes is covered by Dutch law (Wet foetaal weefsel and Article 467 of Behandelingsovereenkomst).

#### Isolation and culture of intestinal crypts

Intestinal crypts containing intestinal stem cells were isolated from whole human foetal small intestine and cultured in 3x 10 μL Matrigel (Corning, 356231) domes per well in a 24-well plate. 500 μL of IntestiCult™ Organoid Growth Medium (Stemcell™ Technologies, 06010), supplemented with 100 U/mL Pen-Strep (ThermoFisher, 15140-122) to maintain enteroid cultures at 37°C, 5% CO2. Every 2-3 days medium was changed and enteroids were passaged twice a week by mechanical dissociation as described previously [22].

Millicell® inserts (Millicell Hanging cell culture insert, PET 3.0 µm, 24-well, Millipore) were coated with 100 µL of 20 µg/mL collagen type I (5 mg/mL, rat tail, IBIDI, 50201) in 0.01% (v/v) acetic acid (VWR, 20103-295) for 1 h at room temperature. Collagen solution was aspirated and inserts were washed twice with PBS (Lonza, 15140-122). Enteroids were then harvested on day 4 after passaging, by enzymatic dissociation using TryplE™ (Gibco, 12605-010) for 10 min at 37°C. Subsequently, 100 μL of a 1*10e6 cells/mL cell suspension was added to the AP side of the coated inserts and 600 μL of IntestiCult™ Organoid Growth Medium, supplemented with 10 μM Y-27632 (Sigma, Y0503), was added to the basolateral (BL) side. After 3 days, medium was changed and fresh medium without Y-27632 was added. After 7 days, medium was changed to differentiation medium containing a 1:1 mixture of IntestiCult™ OGM Human Basal Medium (#100-0190) and Advanced Dulbecco’s Modified Eagle Medium/Nutrient Mixture F-12 (DMEM/F12, ThermoFisher Scientific, 12634-028) supplemented with 1% Pen-Strep, 1% HEPES (Sigma-Aldrich, H3375) and 1% Glutamax (ThermoFisher, 35050-038) to induce differentiation. Medium was refreshed again on day 10 and 13.The monolayers were cultured for 14 days after cell seeding, prior to infection,.

On days 3, 7, 10, and 14 Trans-epithelial electrical resistance (TEER) was measured (EVOM-2 voltohmmeter, World Precision Instruments) 10 min after medium change. Average TEER values were calculated from three measurements per insert. To calculate resistance to surface area (Ω*cm2) the background TEER (empty insert coated with collagen) was subtracted and the result was multiplied by the surface area of the insert (0.3 cm^2^). As a quality control, only inserts with TEER values above 200 Ω*cm2 were used in infection experiments.

### Infection and co-culture with immune cells

Epithelial monolayers were incubated with or without *R. torques* (approx. 10e9 CFU/mL) under anaerobic conditions at 37°C overnight. The next day, AIEC or *R. torques* (approx. 10e9 CFU/mL, multiplicity of infection of 10 (MOI10))) were added to the plate and incubated at 37°C for an additional 6h under anaerobic conditions. Supernatants were harvested for cytokine measurement.

To investigate the crosstalk between the gut epithelial cells and immune cells, PBMCs were incorporated in the CaCo-2 model. Peripheral blood mononuclear cells (PBMCs) from healthy blood-bank donors were isolated by Ficoll-Paque density centrifugation, cryopreserved and stored at - 135°C in 10% dimethyl sulfoxide (DMSO, Sigma Aldrich) and foetal calf serum (FCS) until further use. After thawing, PBMCs (10e6 cells/well) were added to the basolateral compartment of Transwell® inserts on which CaCo-2 cells were cultured apically. Bacterial infection was carried out as described above, and incubated with 5% O2 at 37°C overnight.

Supernatants from both AP and BL compartments were harvested and stored at -20°C for further cytokine measurements, and PBMCs were analysed by flow cytometry.

### Immune profiling by Legendplex and flow cytometry

Immune profiles in supernatants were measured following manufacturer’s instructions with a multiplex bead-based assay, which includes the following cytokines and chemokines: CXCL8 (IL-8), IL-6, IL-10, IL1-beta, IL-18, TNF-alpha, CXCL10 (IP-10), CCL3 (MIP1-alpha), CCL20 (MIP3-alpha), CCL2 (MCP-1), IL-23, IL-33 and IL-12p70 (LEGENDplex™ custom-made Panel (13-plex), BioLegend, San Diego, CA, USA). In brief, standards or supernatants and assay buffer were added to the wells, together with mixed beads and detection antibodies. The plate was subsequently covered and incubated while shaking at 900 RPM for 2h at room temperature, after which Streptavidin-PE (SA-PE) was added to each well. After 30 min incubation, the plate was washed to remove unbound proteins. Subsequently, the samples were acquired by flow cytometry (FACS Canto ll, BD, San Jose, CA, USA) and concentrations were calculated using the LEGENDplex™ Data Analysis Software (BioLegend).

After harvesting supernatants, PBMCs from the basolateral compartment were analysed for immunophenotyping (monocytes, T cells and activation and exhaustion markers) by flow cytometry. Briefly, cells were washed with 0.5% BSA/PBS (PBA) and incubated 15 min with a mixture of antibodies covering the different cell types (see Suppl. Table 1). Samples were acquired on a FACS Symphony (BD) and analysed using FlowJo V10.6.2 Software.

### Migration Assay

Lymphocyte migration was determined by seeding 2x10e6 PBMCs in the AP chamber of a Corning HTS Transwell ® inserts 96-well permeable support with 3.0 μm pore polycarbonate membrane. Two hundred microliters of supernatant from the AP side of the gut epithelial model were added to the BL compartment of the Transwell ® inserts system and the plates were incubated for 2 h at 37°C and 5% CO2. After incubation, cells from the BL compartment were collected and cell type and marker expression was determined with a LSRFortessa X-20 (BD). Data were analysed using FlowJo software version 10.6.2. Count beads were used to determine absolute number of cells.

### Immunofluorescence imaging

Inserts with CaCo-2 cells were fixed in paraformaldehyde (4% v/v) for 15 min at 37°C. Cells were permeabilized with 0.1% (v/v) Triton X-100 for 5 min, blocked with 1% (v/v) BSA, 2% (v/v) Goat-serum (Biolegend) and 0.05% (v/v) Triton X-100 in PBS for 30 min. Subsequently, the cells were stained with Phalloidin (staining actin) and mouse anti-ZO-1 (10 μg/ml, Clone ZO1-1A12, staining tight junctions) for 1 h at 37°C. After washing, goat anti-mouse Alexa Fluor 555 (ZO-1 staining, Biolegend), as a secondary staining, was added for 1 h at 37°C. Nuclei were stained with DAPI (300 nM) for 10 min at 37°C. The polyester membrane containing the cells was excised from the Transwell® insert, placed on a microscopy slide, and embedded in ProLong™ Diamond Antifade Mountant (Invitrogen). Images were captured with a Leica Dmi8 inverted Fluorescence microscope using a Leica DFC7000 GT camera and LAS X 3.4.2 software, with a 40× objective.

Enteroid monolayers were fixed in paraformaldehyde (4% v/v) for 15 min at 37°C and stored in PBS at 4°C. Ice-cold 100% methanol (VWR Chemicals, 20847.240) was added to the inserts for 5 min at room temperature to permeabilize the cells. Methanol was removed and 0.3% (w/v) Sudan Black B (Sigma-Aldrich, 199664-25G) in 70% Ethanol was added for 30 minutes at room temperature. Inserts were removed with a tweezer and washed by immersing the insert into PBS five times.

Subsequently, blocking was performed overnight using SEA BLOCK Blocking buffer (ThermoFisher, 37527) at 4°C. Microscopy slides (VWR, 631-1161) were prepared by circling two parts of the slide with Liquid Blocker Super PAP Pen (Daido Sangyo). Using a scalpel, membranes were cut out from the inserts and placed on the slides within the PAP circle. This was followed by incubation of the cut membranes with primary antibody polyclonal goat anti-human EpCAM (R&D Systems, AF960, 1:100 in SEA BLOCK) overnight at 4°C in a humidified chamber. Membranes were then gently washed three times with Tris-buffered saline (TBS)-Tween (150 mM NaCl, 50 mM Tris-HCL buffer and 1% w/v Tween20) (TBS; EMD Millipore 524750, Tween20; Sigma-Aldrich) and incubated in Alexa Fluor 680 Donkey anti-goat secondary (ThermoFisher, A21084) (1:500 in SEA BLOCK) and Hoechst 33342 (Invitrogen, H3570, 1:1000 in SEA BLOCK) for 2 hours at RT. Membranes were gently washed three times with PBS and mounted using ProLong™ Glass Antifade Mountant (P36984, ThermoFisher Scientific). Slides were imaged using Leica TCS SP8-X microscope with HC Plan Apochromat 40× and 63× oil objective and analysed using Leica LAS X Software (Leica Microsystems).

### Data analysis

All data analyses were performed in R v4.2.0 [28] and RStudio (v2022) with packages ggplot2 [29] and stats [28] among others. Before evaluation of the cytokine data, values were normalized into averages of the logarithms centered by the subtraction of the within-run averages to correct for potential batch effects. Principal component analysis (PCA) was performed using the built-in R function prcomp() and visualized with R packages FactoMineR [30] and factoextra [31]. Pairwise comparisons between the different conditions tested were done using permutational multivariate analysis of variance (PERMANOVA) with the adonis function of the vegan package [32] at 999 permutations.

## Results

### Host response to R. torques and AIEC in a CaCo-2-based intestinal model

We assessed the innate immune response of the CaCo-2 epithelial monolayer to microbial exposure by determining the cytokine and chemokine profiles in the culture supernatants, after stimulation with *R. torques* overnight and, subsequently, challenged with either *R.torques* and AIEC individually or in combination for another 6 h (Suppl. Fig. 1). Principal component analysis (PCA) of the measured immune markers in the apical compartment showed clusters for the different bacterial stimulations (Fig. 1a). Dimension (Dim) 1, explaining 38% of the variation, discriminated clusters of cells infected with the combination of both bacteria or uninfected, while Dim2 (21.9%) mostly separated the conditions stimulated with *R.torques (Rt)* from those stimulated with AIEC. Immune profiles measured differed significantly, except for those observed for stimulations with AIEC and the combination of AIEC and Rt (Fig. 1b). The response of the CaCo-2 monolayer stimulated with Rt was dominated by the production of CXCL8, CXCL10, CCL20 and IL-18, while AIEC induced the production of IL-10, IL-6 and IL-23 (Fig. 1b). When stimulated with the combination of bacteria, levels of cytokine production by individual bacteria were largely maintained, except for IL-18, which was significantly reduced when compared to Rt alone (p= 0.03, Fig. 1b). Under all these conditions, no cytokine or chemokines were detected in the basolateral compartment.

**Fig. 1:**
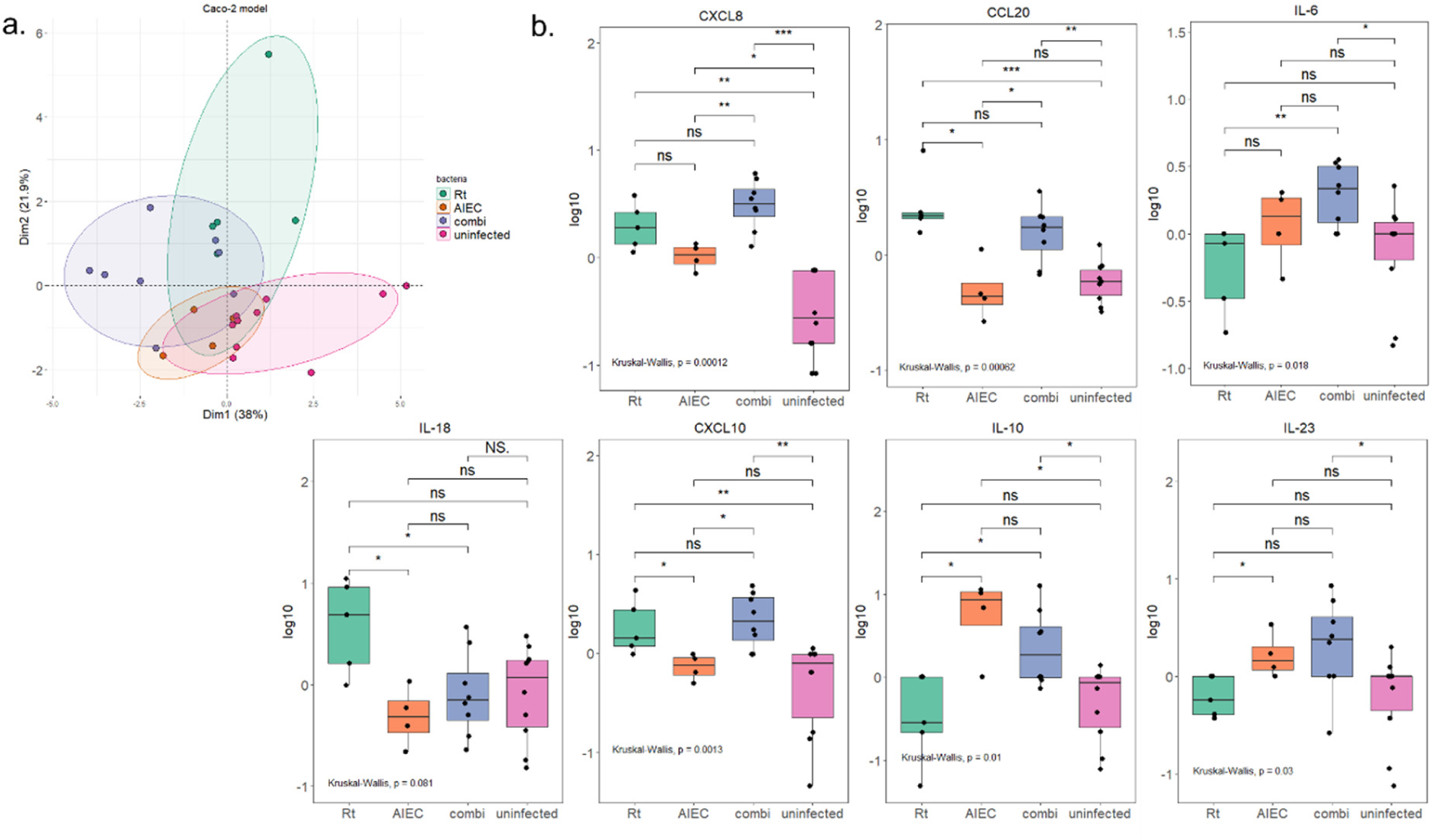
**a) Principal component analysis (PCA) of CaCo-2 cells stimulated with different bacteria individually and in combination.** CaCo-2 cells were challenged with either R.torques and AIEC individually or in combination. PCA was based on the measured cytokines in the apical compartment Rt =Ruminococcus torques (green). AIEC= adherent invasive Escherichia coli (orange). Combi =combination of both bacteria (blue), uninfected (pink). The first two principal axes explained 59.9 % of the variance (Dim = dimension). **b) Cytokine production of CaCo-2 cells stimulated with Rt, AIEC or combination.** CaCo-2 cells produced different cytokines and chemokines upon stimulation with different bacteria. All data were represented as log10 normalized to the average per run for every cytokine or chemokine measured separately. Pairwise comparisons were done using the Wilcoxon test, group comparisons were performed with Kruskal Wallis test. *** p=<0.001 ** p=<0.01 * p=<0.05 ns= p>0.05 NS NS = p=1 .Rt =Ruminococcus torques (green). AIEC= adherent invasive Escherichia coli (orange). Combi =combination of both bacteria (blue), uninfected (pink).

To assess the integrity of the monolayer under these conditions, transepithelial electronical resistance (TEER) was measured throughout the experiment. The epithelial barrier was not disrupted by overnight stimulation with Rt or when Rt or AIEC were added for an additional 6 hours. However, 24h stimulation with Rt or AIEC following the ON stimulation with Rt resulted in barrier loss (Suppl. Fig. 2). Immunofluorescent microscopy was carried out to validate the effect of bacterial stimulation on the tight junctions of the CaCo-2 monolayer. After the 6h infection, cells were stained with ZO-1 (tight junction marker), DAPI (nuclease), and Phalloidin (Actin/cell cytoskeleton). Apical infection with *Rt* showed no impact on the organization of the tight junctions. Conversely, after stimulation with AIEC the epithelial monolayer showed a partial loss of tight junctions and disorganisation, which was less obvious when the infection was performed in combination with *Rt* (Fig. 2).

**Fig. 2:**
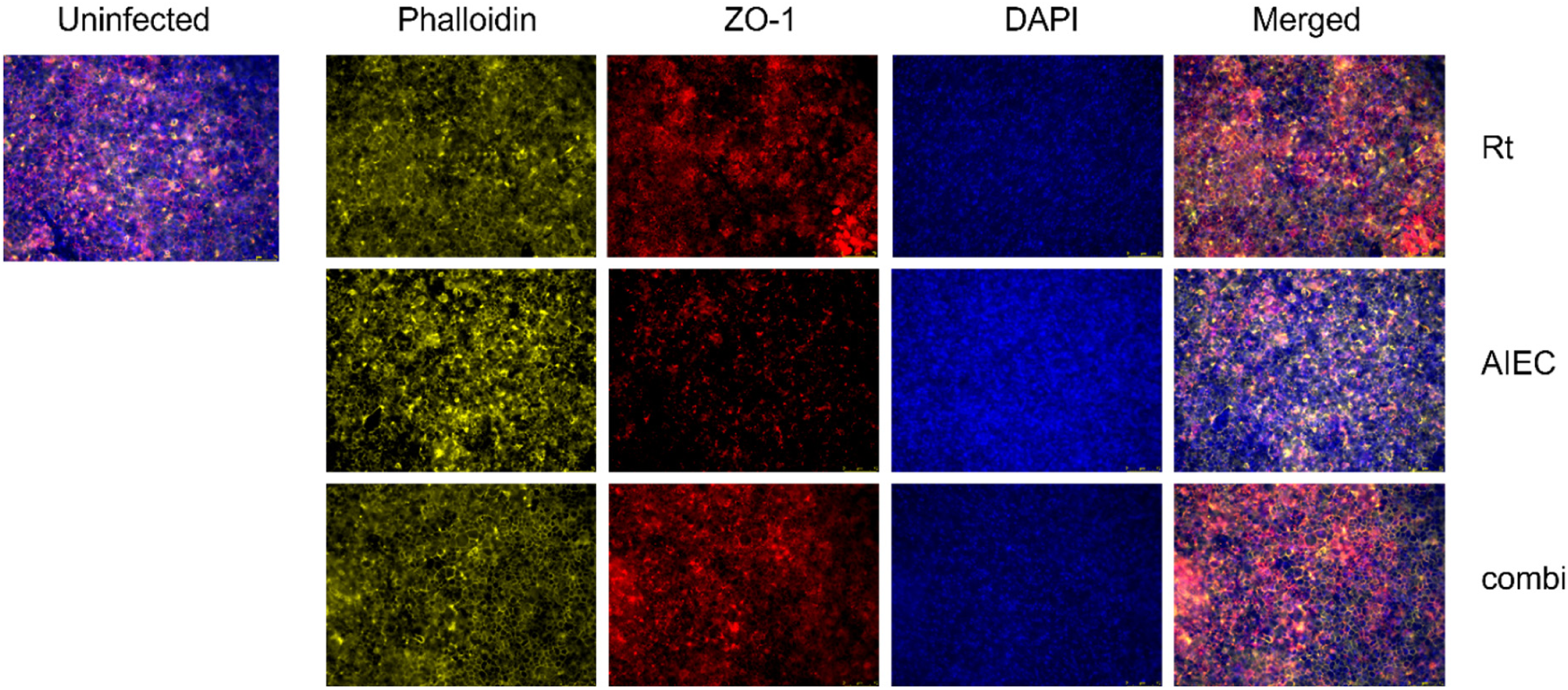
Effect of bacterial stimulation on the permeability of the CaCo-2 monolayer upon stimulation with Rt, AIEC or the combination after 6h. Representative images obtained by fluorescent microscopy of CaCo-2 cultures upon stimulation with the different bacteria, compared to the uninfected monolayer after 6 hours. ZO-1 (tight junction, DSRED), Phalloidin (actin, cell skeleton, Y5), DAPI (nuclease, BLUE)

### Immune activation in co-culture with PBMCs after bacterial stimulation

To investigate how the presence of immune cells in the intestinal ecosystem modulate the epithelial response, PBMCs were included in the basolateral compartment of CaCo-2 cultures during the 6h of infection. The cytokine profile of the CaCo-2 cells measured in the apical compartment, in the presence of PBMC’s in the basolateral compartment, remained comparable to the previously observed responses (Fig. 3a), with higher levels of CXCL8 and IL-6 (Fig. 3b). This suggested that the cytokines detected in the apical compartment were mostly derived from the CaCo-2 cells. When co-cultured with PBMCs, the Rt-induced production of CCL20 was significantly inhibited by addition of AIEC to the overnight stimulation with Rt (Fig. 3b). Although comparable cytokine profiles were observed in the apical compartment regardless of the presence or absence of basolateral PBMCs, principal component analysis (PCA) in co-cultures of CaCo-2 cells and PBMCs (Fig. 3c) revealed distinctive profiles, primarily influenced by Dim2 (23.1%), potentially based on differences of the apical/basolateral polarisation of the intestinal monolayer.

**Fig. 3:**
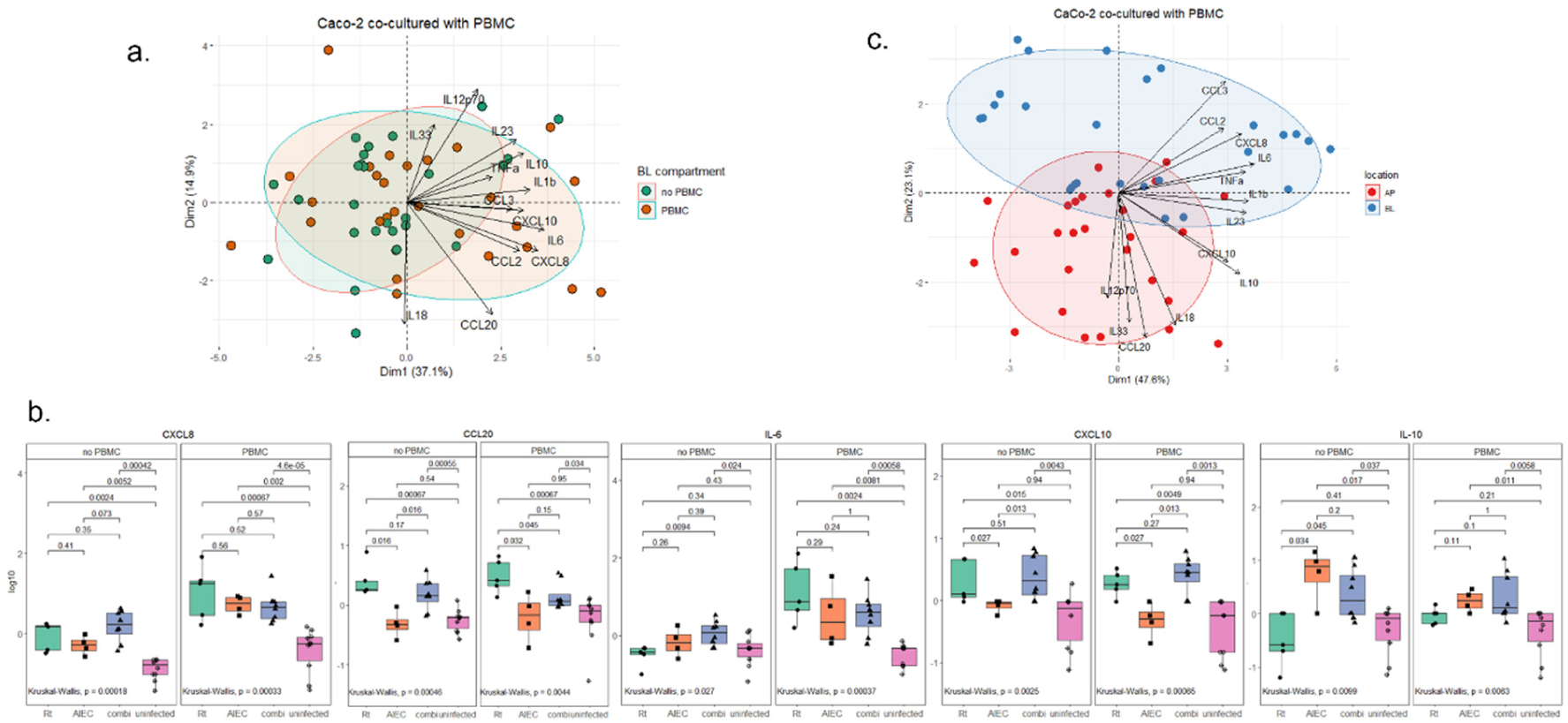
**a) PCA of CaCo-2 cells co-cultured with and without PBMCs in the basolateral (BL) compartment.** Cytokine profiles were measured in the apical compartment, with (orange) or without (green) inclusion of PBMCs in the basolateral compartment. Vectors display the contribution of distinct cytokines. The first two principal axes explained 52 % of the variance (Dim = dimension). **b) Cytokine production in the AP compartment of CaCo-2 cells co-cultured with or without PBMC.** Comparison of the cytokines measured in the apical compartment, with or without PBMC included in the basolateral compartment. Pairwise comparisons were done using the Wilcoxon test, group comparisons were performed with Kruskal Wallis test. Rt =Ruminococcus torques (green). AIEC= adherent invasive Escherichia coli (orange). Combi =combination of both bacteria (blue), uninfected (pink). **c) Biplot of PCA of CaCo-2 cells co-cultured with PBMCs in the basolateral (BL) compartment.** Cytokine profiles were measured in the apical (AP, red) and basolateral (BL, blue) compartments. Vectors display the variables (cytokines measured) contributing to the first two dimensions (70,7%). Dim = dimension.

Closer examination of the different bacterial stimulations in the different compartments showed that responses in the apical compartment were largely uncorrelated with the specific stimulus, as no discernible clusters were observed (Fig. 4). In general, CXCL8, IL-6, CCL2 and IL1-beta were mostly produced after stimulation with Rt, while IL-10, CXCL10 and IL12p70 were produced after AIEC stimulation and the combination of both bacteria. The responses in the basolateral compartment were, however, very distinct between the different bacterial stimulations (Fig. 4). Production of IL-6, IL1-beta and TNF-alpha was observed mainly after stimulation with AIEC and the combination of both bacteria. The responses to the commensal were very similar to the uninfected condition. To investigate the activated fraction of the PBMCs, we performed immunophenotyping by flow cytometry of the basolateral compartment after the different conditions tested on the CaCo-2 model.

**Fig. 4:**
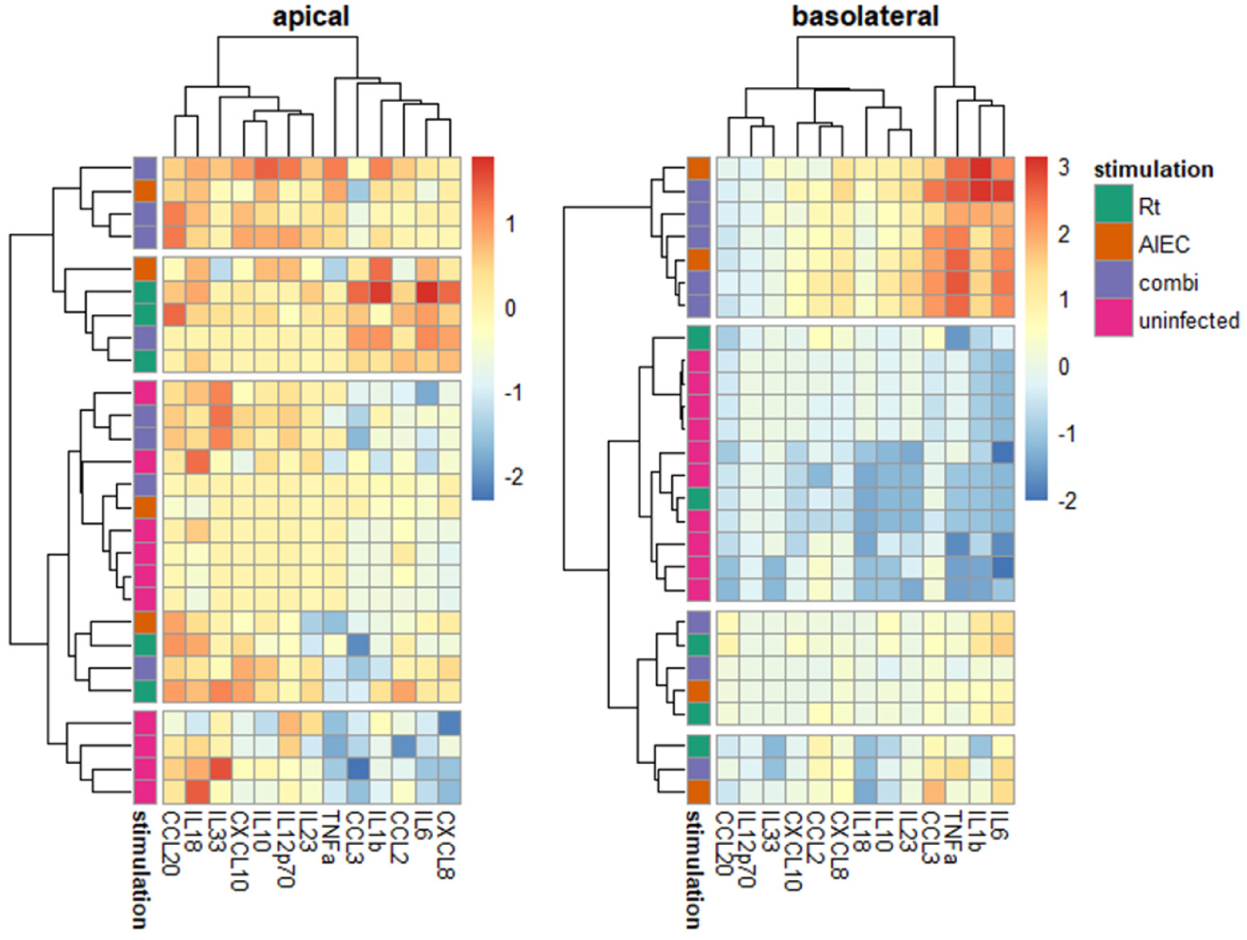
Heatmap of CaCo-2 co-cultured with PBMC. Comparison between the apical (AP) and basolateral (BL) compartment. Distribution of the different cytokines in the apical and basolateral compartment, measured over the different stimulations when the CaCo-2 cells are co-cultured with PBMCs in the basolateral compartment. Rt =Ruminococcus torques (green). AIEC= adherent invasive Escherichia coli.(orange). Combi =combination of both bacteria (blue). uninfected (pink).

As expected, the percentages of the different cell subsets were similar in all conditions studied (Suppl. Fig. S3), stimulation with AIEC showed CD4+ and CD8+ T-cells with a higher activated expression level (CD4+CD69+ and CD8+CD69+) when compared to stimulation with Rt alone. In addition, we used a transmigration assay to investigate which cell subsets were attracted by the chemokines produced by the CaCo-2 cells, using supernatants from the apical compartment in our gut model. Apical supernatants of all bacterial stimulations, particularly Rt, showed migration of different cell subsets (CD4+ and CD8+ T-cells, CD19+ B-cells and monocytes) to the basolateral compartment with PBMCs (Suppl. Fig. S4).

### Host response to R. torques and AIEC in a human intestinal enteroid monolayer model

Apical stimulation of the HIE monolayer with the different bacterial cultures resulted in similar responses compared to CaCo-2 cells. PCA of the measured immune markers revealed that both models showed not only an overlap in their cytokine and chemokine profiles (9 out of 11 cytokines, Fig. 5a), but separation based on the pathogenicity of the bacteria can also be distinguished. While CaCo-2 cells stimulated with the commensal Rt produced significantly higher CCL20, CXCL10 and IL-18, we observed that CCL2 was higher in the HIE monolayer (Fig. 5b). After stimulation with the pathogenic bacteria (AIEC with or without Rt), CaCo-2 cells produced significantly more IL-6 and IL-10, but less IL-18 when compared to the HIE monolayer. Independent of the model, stimulation with Rt showed higher production of CCL20, IL-6, IL-10 and CCL2 than stimulation with AIEC. In general, co-infection did not alter the cytokine production of each individual bacterium in the CaCo-2 model. However, in the HIE model, co-infection led to an increase in CCL20 and a decrease in CCL2 (Suppl. Fig. 5), potentially due to the diversity of cell types present in this model. In addition, despite the use of different donors within the HIE model, clusters based on the pathogenicity are still observed, and likely contribute to the larger variation in the detected immune markers (Fig. 5b and 5c).

**Fig. 5:**
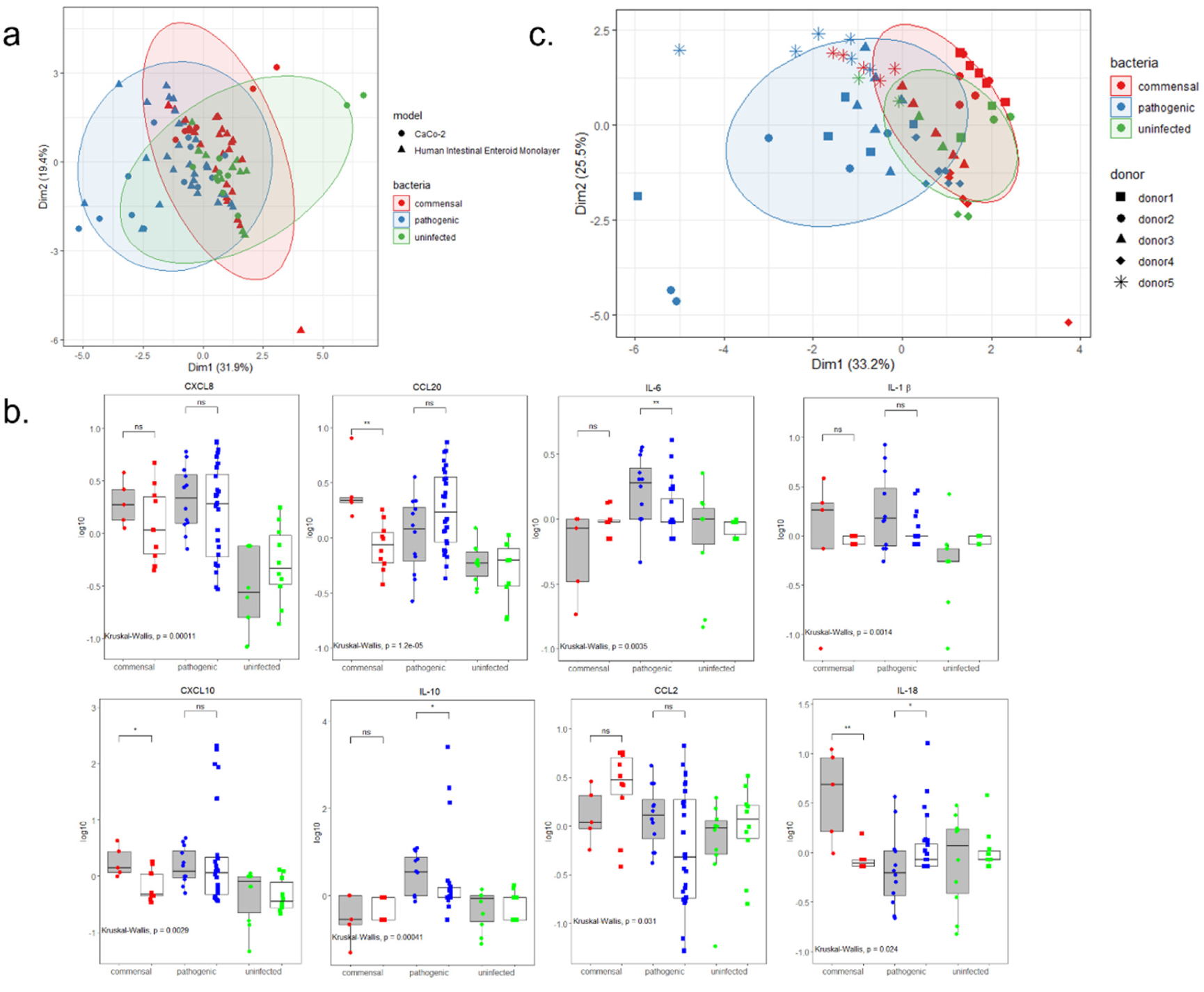
**a) PCA of CaCo-2 monolayer vs. human intestinal enteroid monolayer based on the model and pathogenicity of the bacteria.** Cytokine profiles were measured in the AP compartment in both models after stimulation with different bacteria. Commensal = Rt (red), pathogenic = AIEC and/or in combination with Rt (blue), uninfected (green). **b) cytokine production in apical (AP) compartment in both gut epithelial models.** All data were represented as log10 normalized to the average per run for every cytokine or chemokine measured separately. Pairwise comparisons between the CaCO-2 model (grey bar, circles) and HIE model (white bar, squares) were done using the Kruskal-Wallis test. *** p=<0.001 ** p=<0.01 * p=<0.05 ns= p>0.05 NS NS = p=1. Commensal = Rt (red), pathogenic = AIEC and/or in combination with Rt (blue), uninfected = no infection (green). **c) PCA of human intestinal enteroid monolayer stimulated with different bacteria.** HIE monolayer of 5 different donors (symbols) were challenged with either R.torques and AIEC individually or in combination for another 6 h. PCA was based on the measured cytokines in the apical compartment. Commensal = Rt (red), pathogenic = AIEC and/or in combination with Rt (blue), uninfected (green) Dim=dimension.

Immunofluorescent microscopy of the enteroid monolayer showed that this was partially affected by stimulation with Rt alone, showing gaps in the monolayer (Fig. 6a). However, stimulation with AIEC alone showed clear disruption of the monolayer and lysis of cells (Fig. 6b). In this model in contrast with the CaCo-2 cell based, the monolayer was altered when incubated with Rt overnight, but not as pronounced when compared to the stimulation by AIEC (Fig. 6c).

**Fig. 6:**
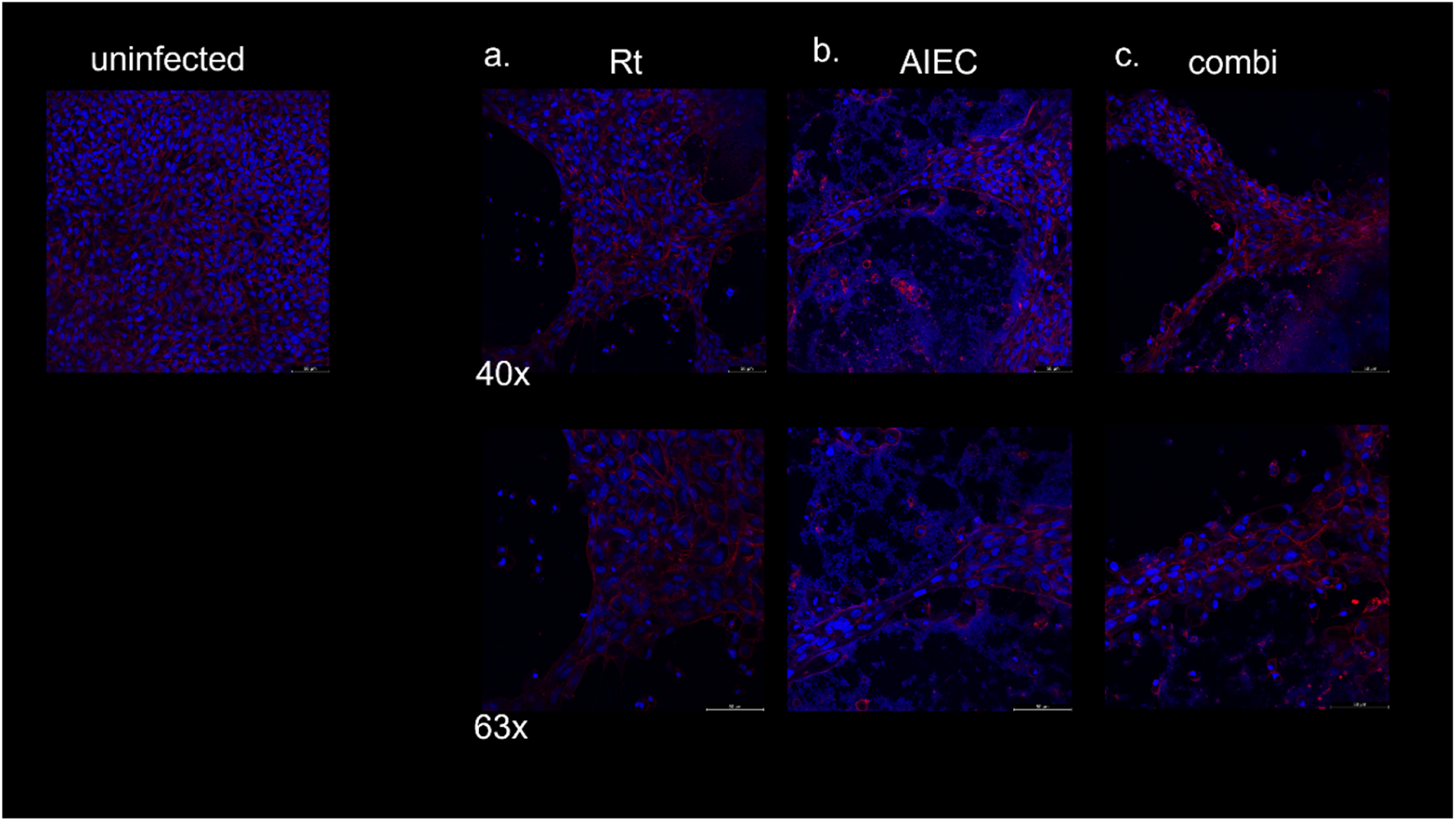
Effect of bacterial stimulation on the permeability of the HIE monolayer upon stimulation with Rt, AIEC or the combination. Representative images obtained by fluorescent microscopy (with 40x or 63x objective) of HIE cultures upon stimulation with the different bacteria, compared to the uninfected monolayer. **a)** Rt =Ruminococcus torques. **b)** AIEC= adherent invasive Escherichia coli. **c)** Combi =combination of both bacteria. Blue = Hoechst (nuclei marker), red = Epcam (epithelial cell marker)

Supernatants from the apical compartment in the human intestinal enteroid model were pooled and used in a transmigration assay to verify which cell subsets were attracted by the chemokines produced. As observed in the CaCo-2 cell based model, apical supernatants of all bacterial stimulations showed migration of different cell subsets (CD4+ and CD8+ T-cells, CD19+ B-cells and monocytes) to the basolateral compartment with PBMCs (n=3). In this model, no significant differences were observed between the commensal and pathogenic stimulations (Suppl. Fig. 6).

## Discussion

The complex interplay between microbes, the host epithelium and immune cells has only partly been elucidated [5, 6]. In our study, we aimed to investigate the immune processes that occur in the gut epithelium using both a traditional CaCo-2 cell-based epithelial model, and a more representative human intestinal enteroid model. Based on previous observations of a positive association between pro-inflammatory profiles and the presence of *E. coli* and *R. torques* in the gut ecosystem of individuals during an influenza-like illness (ILI) [14], we investigated how these bacteria modulated intestinal epithelial cell responses. Challenging of the epithelial monolayers with the commensal (*R. torques*) and pathogenic (adherent invasive *E. coli*, AIEC) bacteria revealed that we can investigate the pathogenicity of these bacteria, with an overlap in cytokine and chemokine profiles between the models. The bacteria induced different cytokine responses by the epithelial monolayers, and a higher cytopathology was observed for AIEC. Addition of PBMCs in the basolateral compartment of the CaCo-2 system during infection showed that the epithelial responses in the apical compartment are not modulated.

Several studies have shown that members of the *Ruminococcus* genus are associated with local inflammation-mediated disorders, such as IBD [33, 34], but also with inflammatory disorders at distant sites [35, 36]. *R. torques* belongs to a group of commensals known to be associated with chronic gut inflammation [33]. In this study, we demonstrated that stimulation with *R. torques* alone induced pro-inflammatory responses by the epithelial cells, dominated by the production of CXCL8, CXCL10, CCL3 and IL-18. This proinflammatory profile was also observed during influenza-like illness, as shown by our previous study [14]. Also, these Rt-induced pro-inflammatory chemokines are involved in leukocyte and monocyte recruitment, which was corroborated by the transmigration assay with PBMCs. The innate immune system, as a first line in host defence, plays an important role in orchestrating downstream immune responses to pathogens, that could contribute to inflammation and ILI-like complaints [37, 38]. In addition, exacerbated immune responses, besides direct epithelial damage by microorganisms, can also damage the epithelial barrier [3].

The AIEC strain LF82, a strain generally associated with IBD and a plausible driver of intestinal inflammation [11-13], has the capacity to adhere and invade the gut epithelial monolayer. AIEC can trigger signal transduction pathways, which could result in suboptimal epithelial control of invading pathogens [12, 13, 39]. In our study, stimulation of the epithelial monolayers with AIEC showed a distinct cytokine profile that differed from that of *R. torques.* While in the CaCo-2 model the profile induced by AIEC was mainly driven by IL-10, IL-6 and IL-23 production, in the HIE model CXCL10 and CCL20 were also produced. Co-infection did not impact the cytokine production induced by the commensal, although IL-18 was significantly reduced by AIEC, suggesting an active inhibition of at least this particular component of the immune response. This suppressed response could be a result of a damaged epithelium followed by infection, leading to inflammation and potential barrier loss with the result of direct exposure of bacteria to immune cells. Local inflammasome-mediated IL-18 plays an important role in the regulation of homeostasis in the gut [40-44], and is induced upon infection with influenza virus with a role in the antiviral activity of the immune system.

We further investigated how *R. torques* and AIEC may impact downstream host immune responses in the gut epithelium. Addition of PBMCs to the basolateral compartment did not impact the cytokine and chemokine production of the CaCo-2 cells at the apical side. In fact, cytokine responses upon stimulation with Rt resembles the uninfected epithelium, suggesting a homeostatic response where PBMCs do not trigger additional responses in the CaCo-2 cells to the bacterial stimulations. On the basolateral compartment, while infection did not lead to production of immune markers in the absence of PBMCs, addition of these cells showed a proinflammatory cytokine profile mainly induced by infection with AIEC.

Despite the fact that the CaCo-2 cell-based epithelial model has been shown in ours and other studies as a useful tool to model infection and host responses, it has limitations due to the lack of cellular diversity. Recently developed human primary intestinal organoid models can alleviate some of these issues. Although these models are costly and technically challenging [25], they provide more insights into host-pathogen interactions comparable to the in vivo situation [22, 45, 46]. In our study, comparison of the immune responses induced by the different bacteria in both models showed similar cytokine and chemokine profiles. In fact, stimulation with commensal and pathogenic bacteria can be clearly differentiated independent of the model. Stimulation of HIE with the pathogenic bacteria showed not only low production of CCL2, a member of chemotactic cytokines which attracts macrophages and also enhances pro-inflammatory responses [47, 48], but also inhibition in co-infection. In line with this, we observed higher production of IL-6 and IL-10 induced by stimulation with the pathogenic bacteria in both gut epithelial models. In addition, it has been established that *R. torques* can degrade the mucus layer by colonizing the intestinal mucosa [33], and release of mucins and other substrates from the mucus layer, serve as a substrate beneficial for AIEC to overgrow in the gut [49]. The main drawback of our CaCo-2 model is the fact that Caco-2 cells do not produce significant amounts of mucins under normal growth conditions [50]. The human intestinal enteroid monolayer model, on the contrary, contain all cell types of the intestinal epithelium present in vivo, including the MUC2 expressing goblet cells [22]. Fluorescent imaging of the human intestinal enteroid monolayer supports the hypothesis that overnight priming of the monolayer with *R. torques* is beneficial for invasion of AIEC, as gaps in the monolayer become visible.

Taken together, we demonstrate that both CaCo-2 cell-based model and a human intestinal enteroid (HIE) model are suitable to understand the immune processes that occur in the gut epithelium in the presence of commensal and pathogenic bacteria, nevertheless both models have their own limitations. While the CaCo-2 model represents a simple cell culture model, it lacks the complexity to study the mechanisms of interaction between the commensal and pathogenic bacteria. The human intestinal enteroid model represents the epithelial cell composition in vivo more accurately, but lacks the dynamics of the intestinal mucosa. Although the human enteroid model should be further examined on host immune responses, this model could be a more promising system and valuable tool to the complexity of gut-host-microbiome interactions. Moreover, increasing the complexity of the microorganisms added, could give more insights into the immunomodulatory effects of bacteria on the gut ecosystem.

## Supplementary Material

**Supplementary Table 1:**
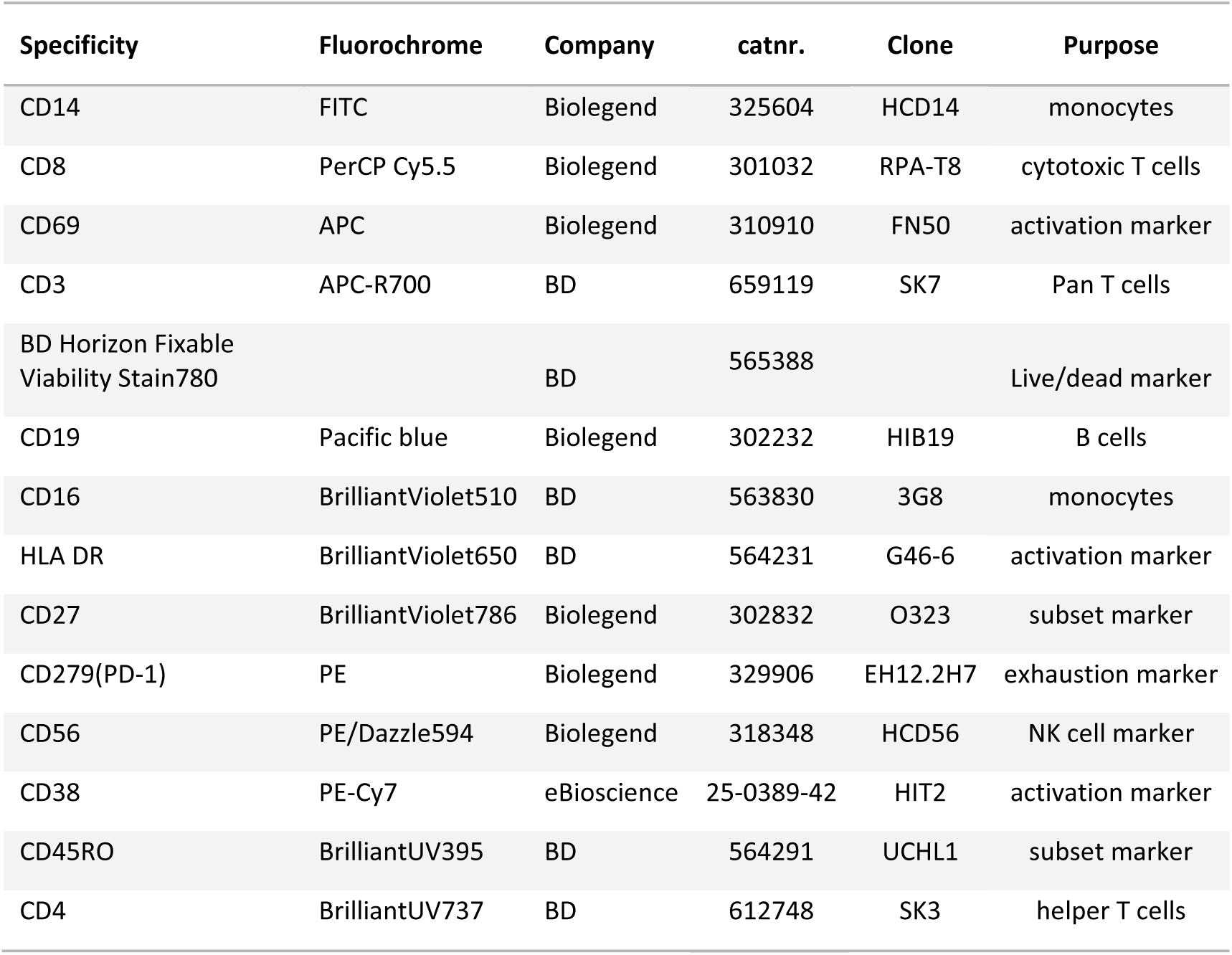
Antibodies used for Flow Cytometry.

| Specificity | Fluorochrome | Company | catnr. | Clone | Purpose |
| --- | --- | --- | --- | --- | --- |
| CD14 | FITC | Biolegend | 325604 | HCD14 | monocytes |
| CD8 | PerCP Cy5.5 | Biolegend | 301032 | RPA-T8 | cytotoxic T cells |
| CD69 | APC | Biolegend | 310910 | FN50 | activation marker |
| CD3 | APC-R700 | BD | 659119 | SK7 | Pan T cells |
| BD Horizon Fixable Viability Stain780 |  | BD | 565388 |  | Live/dead marker |
| CD19 | Pacific blue | Biolegend | 302232 | HIB19 | B cells |
| CD16 | BrilliantViolet510 | BD | 563830 | 3G8 | monocytes |
| HLA DR | BrilliantViolet650 | BD | 564231 | G46-6 | activation marker |
| CD27 | BrilliantViolet786 | Biolegend | 302832 | O323 | subset marker |
| CD279(PD-1) | PE | Biolegend | 329906 | EH12.2H7 | exhaustion marker |
| CD56 | PE/Dazzle594 | Biolegend | 318348 | HCD56 | NK cell marker |
| CD38 | PE-Cy7 | eBioscience | 25-0389-42 | HIT2 | activation marker |
| CD45RO | BrilliantUV395 | BD | 564291 | UCHL1 | subset marker |
| CD4 | BrilliantUV737 | BD | 612748 | SK3 | helper T cells |

## Figure Legends

**Supplementary Figure 1:**
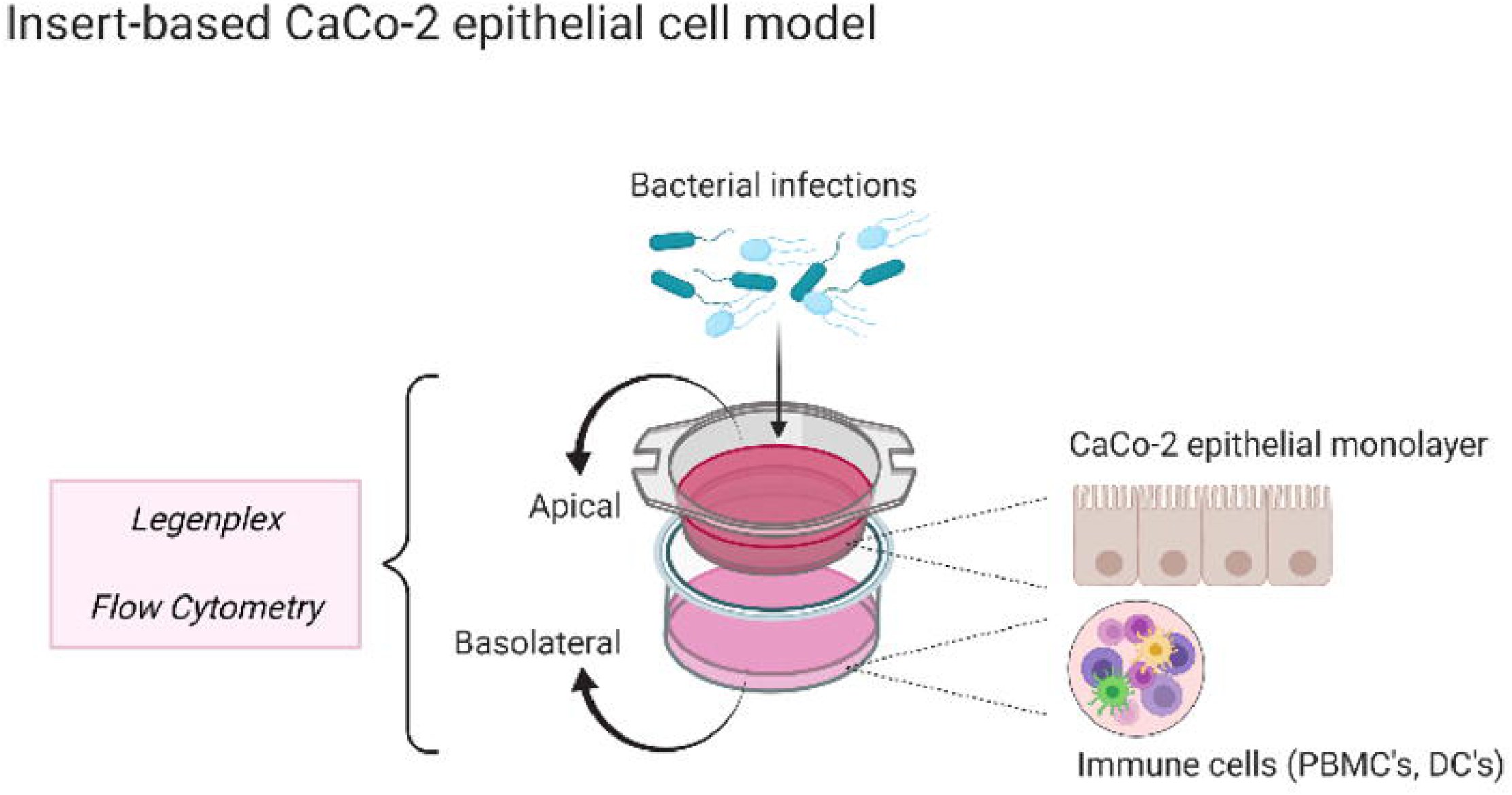
Schematic illustration of the insert-based CaCo-2 epithelial cell model. The CaCo-2 monolayer, cultured at the apical side of the insert, was stimulated with R.torques (RT) overnight and subsequently challenged with RT and/or AIEC for another 6 h. Culture supernatants were harvested to measure cytokines and chemokines. For co-cultures, Immune cells were included in the basolateral compartment for downstream analysis.

**Supplementary Figure 2:**
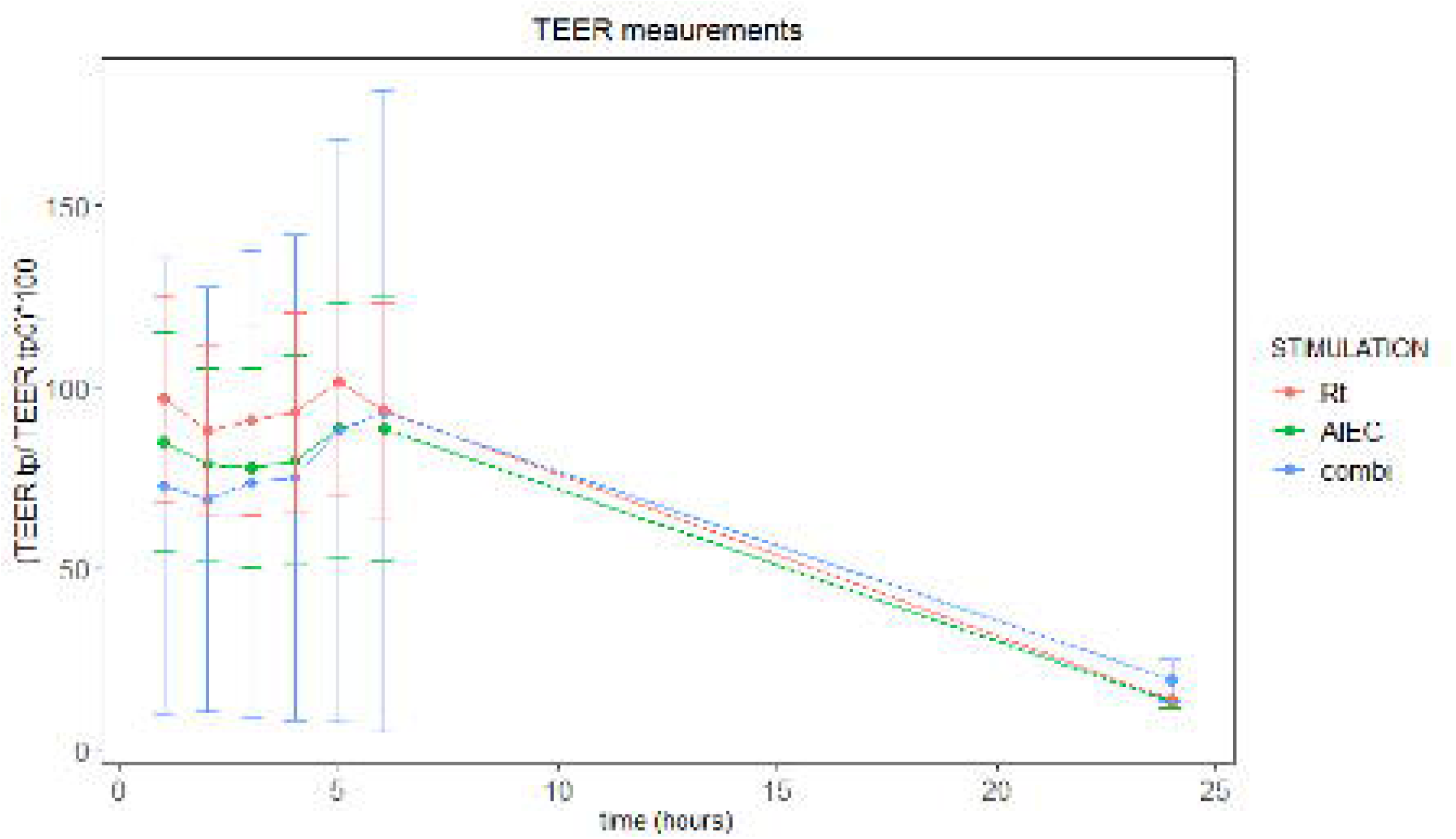
Effect of bacterial stimulation on the permeability of the CaCo-2 monolayer upon stimulation with Rt, AIEC or the combination. TEER values are presented as relative variations with respect to the unstimulated condition. These results are means ± SEM for eight measurements for all conditions tested. tp = timepoint, tp0 = timepoint 0

**Supplementary Figure 3:**
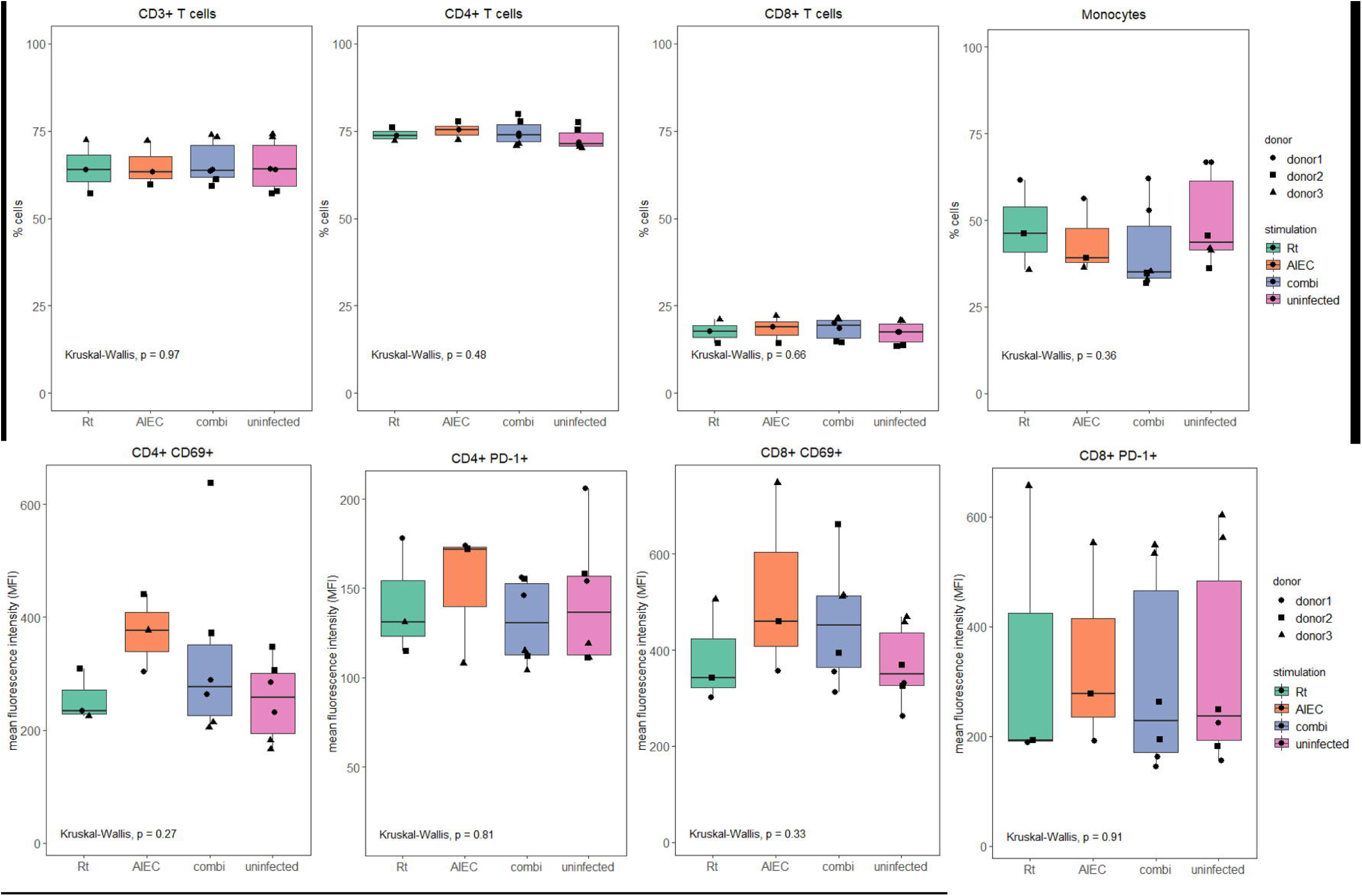
Phenotyping of the PBMCs in the BL compartment after different co-culturing conditions of the intestinal epithelial model. PBMCs of 3 different donors (symbols) were stained for immunophenotyping (monocytes, T cells and activation (expressed as percentages) and exhaustion markers (expressed as mean fluorescence intensity)) and analyzed by Flow Cytometry. Pairwise comparisons were done using the Wilcoxon test, group comparisons were performed with Kruskal Wallis test. *** p=<0.001 ** p=<0.01 * p=<0.05 ns= p>0.05 NS NS = p=1 .Rt =Ruminococcus torques (green). AIEC= adherent invasive Escherichia coli.(orange). Combi =combination of both bacteria (blue). uninfected (pink).

**Supplementary Figure 4:**
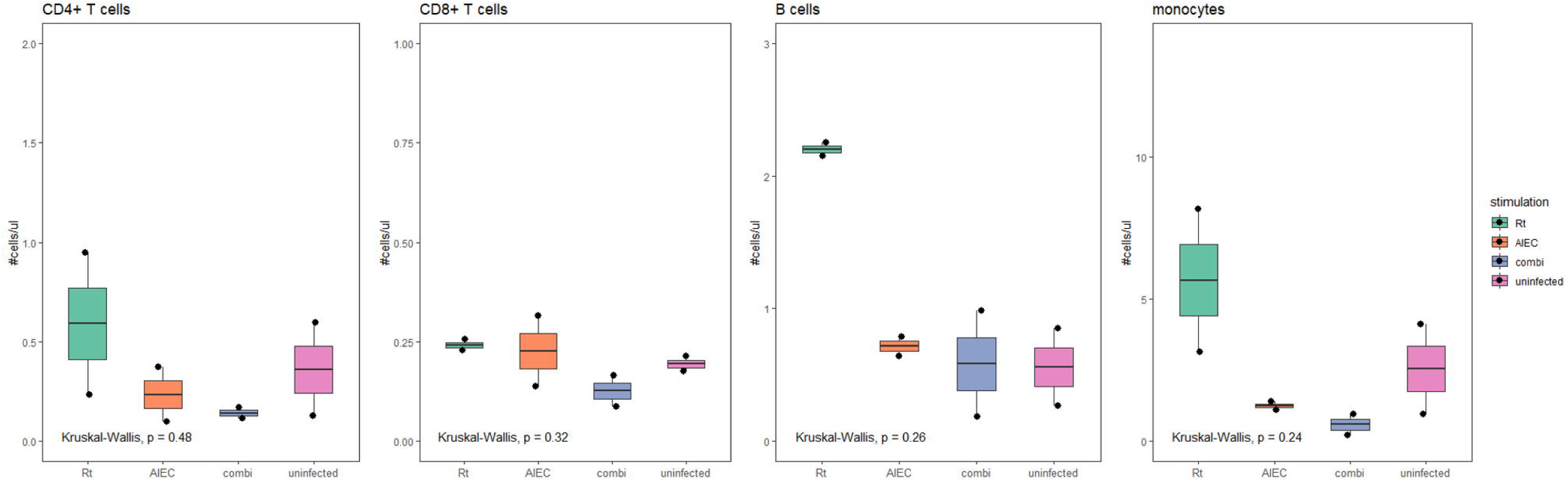
Lymphocyte migration assay within the CaCo-2 model. Supernatants of the AP compartment of the CaCo-2 model were used to perform a transmigration assay. Pairwise comparisons were done using the Wilcoxon test, group comparisons were performed with Kruskal Wallis test. *** p=<0.001 ** p=<0.01 * p=<0.05 ns= p>0.05 NS NS = p=1 Rt =Ruminococcus torques (green). AIEC= adherent invasive Escherichia coli.(orange). Combi =combination of both bacteria (blue). uninfected (pink).

**Supplementary Figure 5:**
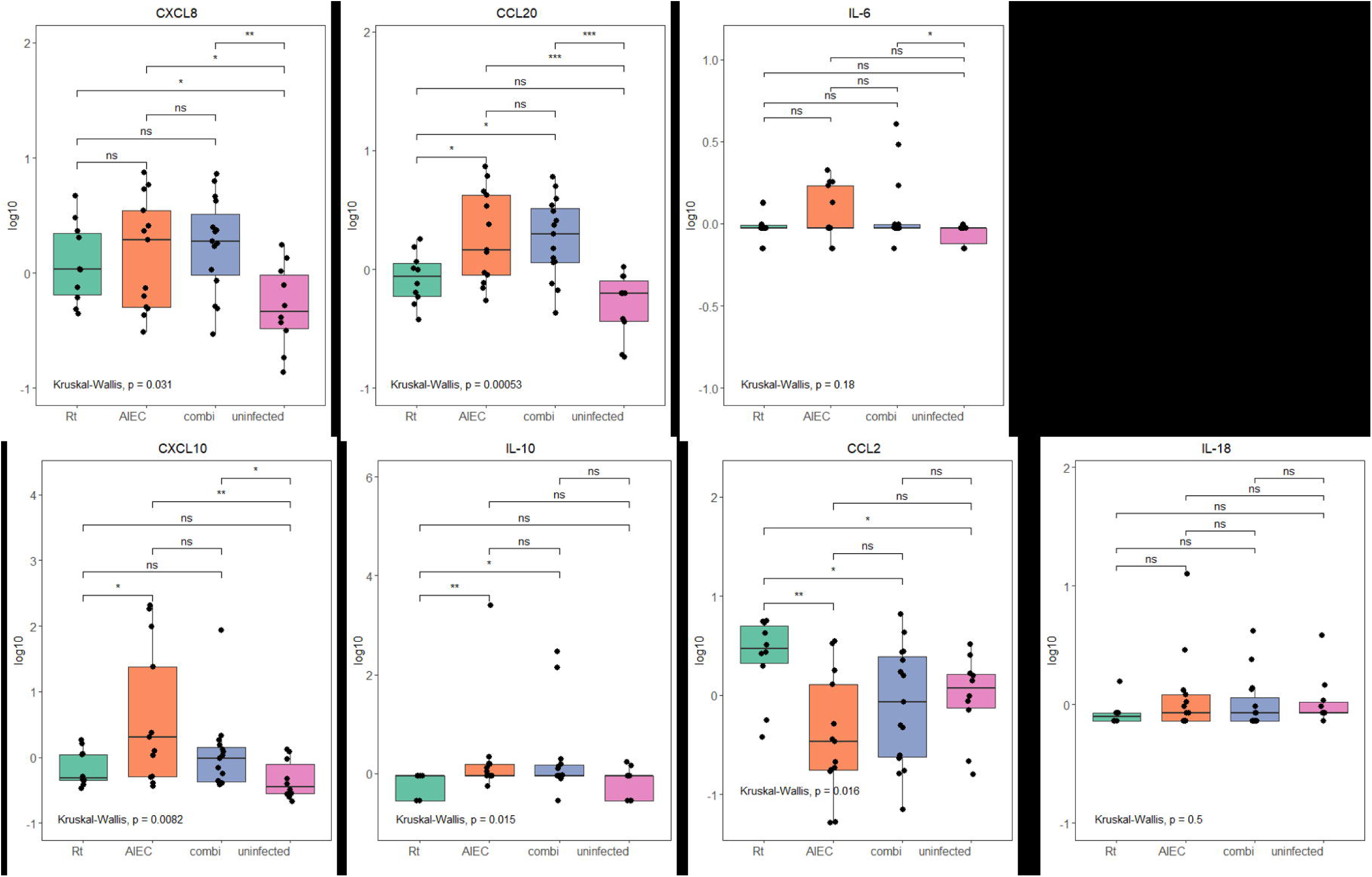
Cytokine production of HIE monolayer stimulated with Rt, AIEC or combination. HIE monolayer produced different cytokines and chemokines upon stimulation with different bacteria. All data were represented as log10 normalized to the average per run for every cytokine or chemokine measured separately. Pairwise comparisons were done using the Wilcoxon test, group comparisons were performed with Kruskal Wallis test. *** p=<0.001 ** p=<0.01 * p=<0.05 ns= p>0.05 NS NS = p=1. Rt =Ruminococcus torques (green). AIEC= adherent invasive Escherichia coli (orange). Combi =combination of both bacteria (blue), uninfected (pink).

**Supplementary Figure 6:**
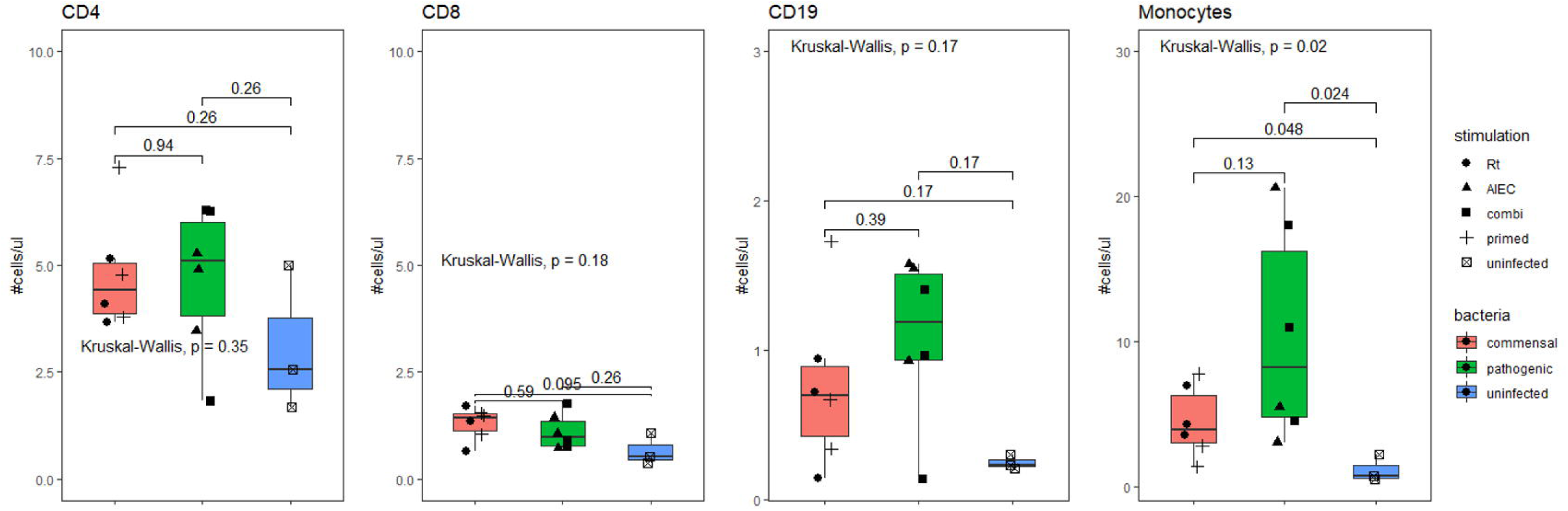
Lymphocyte migration assay within the human intestinal enteroid monolayer. Supernatants of the AP compartment of the HIE model were used to perform a transmigration assay. Pairwise comparisons were done using the Kruskal-Wallis test. Rt =Ruminococcus torques. AIEC= adherent invasive Escherichia coli. Combi =combination of both bacteria. Commensal = Rt (red), pathogenic = AIEC and/or in combination with Rt (blue), uninfected (green)

